# Progressive Supranuclear Palsy-associated PERK haplotype B selectively enhances DLX1 translation and promotes tau toxicity

**DOI:** 10.1101/2025.06.30.662315

**Authors:** Christian Lessard, Diego Rubio Rubio, Samantha Tolton, Marangelie Criado-Marrero, Sakthivel Ravi, Tristyn N. Garza, John Koren, Jennifer Philips, Pritha Bagchi, Karen McFarland, Deepak Chhangani, Todd Golde, Benoit I. Giasson, Jada Lewis, Paramita Chakrabarty, Matthew J. LaVoie, David Borchelt, Nicholas T. Seyfried, Stefan Prokop, Diego E. Rincon-Limas, Jose F. Abisambra

## Abstract

The unfolded protein response (UPR) sensor PERK exists in two haplotypes termed A and B. PERK-B uniquely confers increased risk for tauopathies like progressive supranuclear palsy (PSP), but the mechanisms distinguishing its function from PERK-A and contributing to its association with tau pathology are not known. Here, we developed a controlled cellular model for a pair-wise comparison of the two PERK haplotypes, finding their UPR functions nearly indistinguishable. However, a careful examination employing puromycin-based proteomics revealed that a subset of mRNA translation events were permissible under PERK-B, but not PERK-A, dependent UPR. Critically, one of the targets that escaped PERK-B suppression was the transcription factor DLX1, which has been genetically linked to PSP risk. Here, we found a shift in the solubility of DLX1 in human PSP brain tissue, and report that silencing of DLX-1 reduced the aggregation of tau in mammalian cells. Furthermore, silencing of the fly homolog of DLX1 was sufficient to decrease tau-induced toxicity, *in vivo*. Our results detail the haplotype-specific PERK-B/DLX-1 pathway as a novel driver of tau pathology in cells, flies, and likely human brain, revealing new insights into PSP pathogenesis and potential therapeutic targets.

## Introduction

Cells undergoing endoplasmic reticulum (ER) stress activate an adaptive tripartite mechanism called the unfolded protein response (UPR). The protein kinase ER-like kinase (PERK) pathway of the UPR attenuates translation by phosphorylating and thereby inhibiting the eukaryotic initiation factor 2α (eIF2α) ^1,2^. While phosphorylated eIF2α broadly suppresses translation, it selectively permits translation of transcription factors such as ATF4, which promotes expression of proteins that restore ER function. Therefore, in response to ER stress, PERK regulates both transcription and translation ^3,4^. Failure to resolve ER stress, accompanied by long-term PERK activation, can lead to broad and sustained inhibition of protein synthesis and ultimately, cell death ^5^.

Given that neuronal function requires *de novo* protein synthesis, long-term attenuation of translation is problematic, especially for neurons ^6,7^. Maladaptive PERK activity leads to sustained suppression of translation and deleterious brain health. For example, PERK haploinsufficiency results in Wolcott-Rallison Syndrome, which features developmental brain abnormalities ^8^. Consistent with the pathologic effects of PERK deficiency, expression of the PERK haplotype B (PERK-B) produces a hypomorphic PERK variant compared to haplotype A (PERK-A), and it is associated with higher risk of developing tauopathies such as progressive supranuclear palsy (PSP) and Alzheimer’s disease (AD) ^9–13^. However, the mechanisms by which PERK-B contributes to tauopathy remains unknown.

Besides the genetic link, PERK is also implicated in tauopathies. Pathologically, active PERK (phosphorylated, or ‘pPERK’) accumulates in pre-tangle neurons of tauopathy brains, including PSP, AD, Down Syndrome, and frontotemporal dementia (FTD)-tau ^14–18^. Moreover, accumulation of pathological tau drives chronic PERK activation without resolution of ER stress, further hampering neuronal function^19^. While modifying PERK activity ameliorates accumulation of pathological tau, whether activating or inhibiting PERK confers benefits remains unclear ^7,9,12,20,21^.

To define the enigmatic mechanism by which PERK drives diverging cell fates, we developed novel tools to specifically activate PERK-A and PERK-B and found no differences in their canonical functions. Using puromycin proteomics, we established translatomes of both PERK haplotypes, and the overwhelming number of translational targets were equally suppressed by both PERK-A and PERK-B. Critically, the expression of a small subset of proteins was uniquely preserved under PERK-B-induced ER stress that was not present with PERK-A. Among these novel targets was DLX1, a gene product strongly associated with PSP ^22^. These data reveal a novel pathway linking PERK function and genetic risk of PSP with mechanisms underlying the deposition of tau in cellular and Drosophila models of disease.

## Results

### PERK overexpression model

To examine the impact of PERK haplotypes A and B individually and independently of endogenous PERK expression, we generated a PERK knockout human embryonic kidney (PERK-KO HEK) cell line using a CRISPR/Cas9 strategy. We also produced plasmid constructs of PERK-A, PERK -B, and PERK-K (an enzymatically dead PERK variant, K622R, used as a negative control). Experimentally, ER stress is commonly induced with chemical compounds (*e.g.*, thapsigargin and tunicamycin) that promote *en masse* calcium efflux from the ER ^23^. To bypass confounds of chemically induced excitotoxicity, we transfected cells with varying amounts of plasmids expressing PERK-A, PERK -B, or PERK-K in PERK-KO HEK cells for 48 hours (Fig. 1A). We found that transient transfection with 100ng of PERK variants was not toxic and induced PERK activation (Fig. 1A-B).

**Figure 1.**
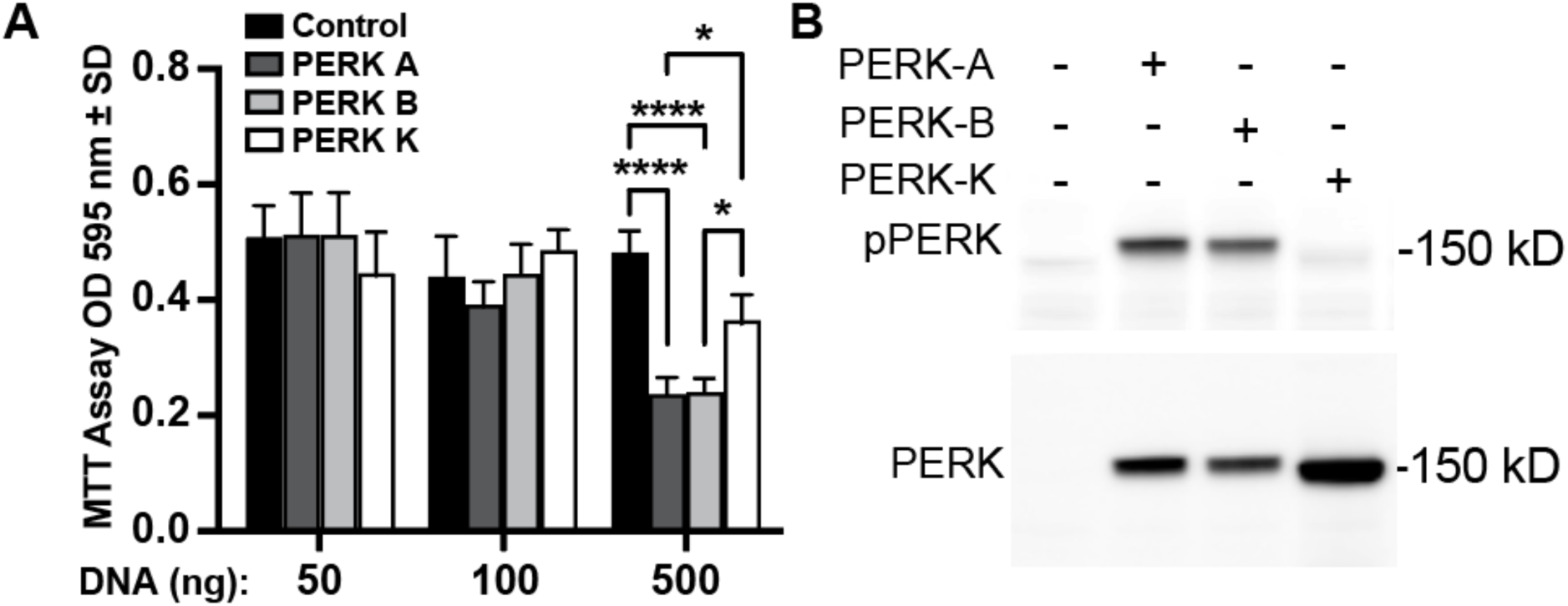
Mild PERK-A and PERK-B overexpression preserves cell viability while activating PERK. PERK knockout (KO) HEK293T cells were transfected with varying low (nanogram) amounts of mammalian expression plasmids encoding PERK-A, PERK-B, PERK-K or an empty plasmid used as a carrier control. (A) Cell viability was assessed 48 hours post-transfection using an MTT assay. No significant cellular toxicity was detected at 50 and 100 ng of transfected PERK plasmids. Results were reported as the OD595 averaged ± standard error (n=3; * p<0.05, **** p < 0.0001, Two-way ANOVA, Turkey’s multiple comparisons). (B) Western blot analysis confirming that PERK and phosphorylated PERK were detectable when cells were transfected with 100 ng of transfected plasmid. The K622R mutation in PERK did not lead to phosphorylation validating the specificity of PERK activation.

### PERK haplotypes share common pathways and have similar effects on translation

We used ATF4 and Xbp1 sensors to determine the effect of PERK haplotypes on the PERK and Ire1 branches of the UPR in real time (Fig. 2A). ATF4-sensor fluorescence increased when either PERK-A or PERK-B were expressed (Fig. 2B); as expected, PERK-K failed to induce ATF4 fluorescence, and chemical activation of the UPR with thapsigargin induced ATF4 (Fig. 2B). PERK-A and -B expression also activated the Ire1 pathway as evidenced by increased Xbp1 sensor fluorescence (Fig. 2C); interestingly, PERK-K also activated Xbp1, suggesting that PERK overexpression induces Ire1 independently of pPERK upregulation (Fig. 2C).

**Figure 2.**
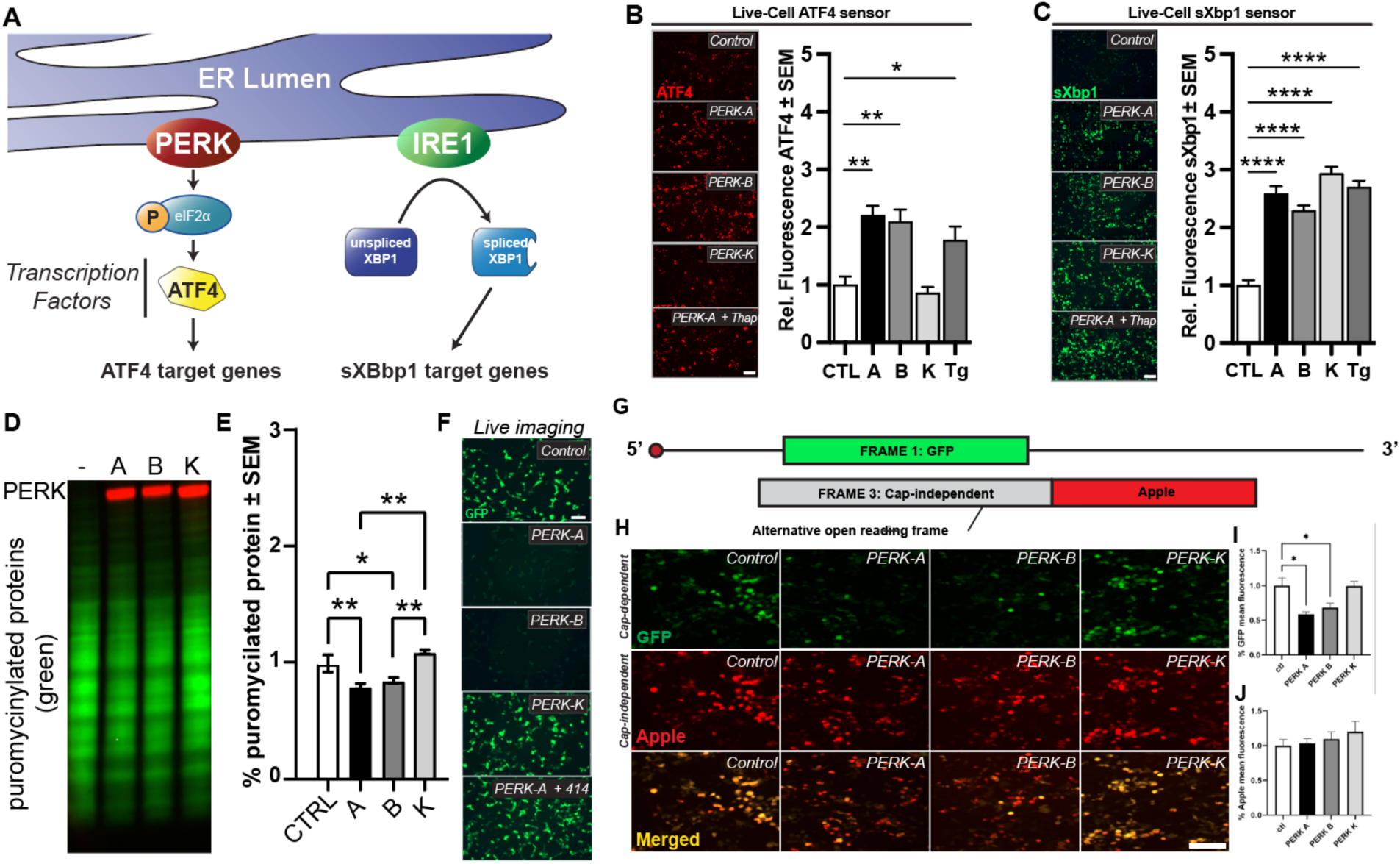
PERK-A and PERK-B canonical functions are indistinguishable. (A) Schematic representation of PERK and Ire1 activation leading to eIF2α phosphorylation and XBP1 mRNA splicing, key events in the unfolded protein response (UPR). (B, C) PERK KO HEK293T cells were co-transfected with 100 ng of either an ATF4-mScarlet sensor (B) or an XBP1-mNeogreen sensor (C) along with 100 ng of PERK (PERK-A, PERK-B, or PERK-K) or an empty plasmid as a carrier. UPR activation was assessed by measuring fluorescence from the ATF4 and XBP1 sensors upon PERK expression or with thapsigargin (Tg) treatment. PERK-K expression also activated the XBP1 sensor. Relative fluorescence quantification shows significantly higher fluorescence levels for the activated sensors compared to controls (n = 3; *p < 0.05, **p < 0.01, ****p < 0.0001, one-way ANOVA, Dunnett’s multiple comparisons). (D, E) PERK KO HEK293T cells transfected with 100 ng of PERK constructs plus an empty plasmid were incubated with puromycin for 1 hour. Western blot analysis shows reduced levels of puromycinylated proteins in cells expressing PERK-A or PERK-B, indicating repression of protein synthesis. Quantification of puromycinylated proteins is presented as mean ± SEM (n = 3; *p < 0.05, **p < 0.01, one-way ANOVA, Tukey’s multiple comparisons). (F) Live imaging of PERK KO HEK293T cells transfected with 100 ng of PERK constructs and a GFP-expressing plasmid. Expression of PERK-A or PERK-B significantly reduced GFP fluorescence. The addition of a PERK inhibitor post-transfection reversed the PERK-A effect on GFP fluorescence. (G) Schematic representation of the GFP-Apple construct, where GFP is expressed in frame 1 (cap-dependent) and Apple is expressed in frame +3 (cap-independent) following the GFP stop codon. This design allows visualization of cap-independent translation through Apple fluorescence. (H) Live imaging of PERK KO HEK293T cells transfected with 100 ng of PERK constructs and the GFP-Apple plasmid. PERK-A and PERK-B expression reduced GFP fluorescence while allowing detectable Apple fluorescence, demonstrating selective repression of cap-dependent translation. (I, J) Quantification of GFP fluorescence shows a significant reduction with PERK-A and PERK-B expression, while Apple fluorescence remains unaffected.

Given that active PERK levels were detectable in PERK-A and -B overexpressing cells, we hypothesized that both PERK variants suppressed translation. To test this, we performed surface sensing of translation (SUnSET), a method that uses puromycin to label nascent proteins ^24^. PERK-A and -B expression reduced puromycinylated proteins, and the decrease was similar in both conditions (Fig. 2D-E). Furthermore, we performed live-cell imaging and found that GFP levels, which served as a surrogate of protein synthesis, were significantly reduced by PERK-A and -B, but not -K; likewise, both PERK-A and -B equally reduced GFP signal (Fig. 2F). PERK-A overexpression in combination with a PERK inhibitor (GSK2606414) rescued GFP fluorescence (Fig. 2F).

PERK reduces cap-dependent translation ^25^. To determine whether PERK-B reduced translation following the same mechanism, we generated a construct that measures cap-dependent translation by GFP expression; using the same construct in a different frame produces red (apple) fluorescence, which represents cap-independent translation (Fig. 2G). Similar to PERK-A, PERK-B reduced translation in a cap-dependent manner (Fig. 2H-J). As expected, neither construct reduced cap-independent translation.

### PERK haplotypes differentially impact the levels and aggregation of pathological tau species

Given that PERK-B confers increased risk for tauopathies, we compared the effects of PERK-A and PERK-B on tau stability. We created a dose- and time-dependent window of PERK expression by transfecting 100ng of PERK expression plasmids for 16hrs. In this acute time frame, both PERK-A and -B induced eIF2α phosphorylation and reduced PHF1 (pS396/S404) and total tau levels compared to both the empty vector and PERK-K controls (Fig. 3A-C).

**Figure 3.**
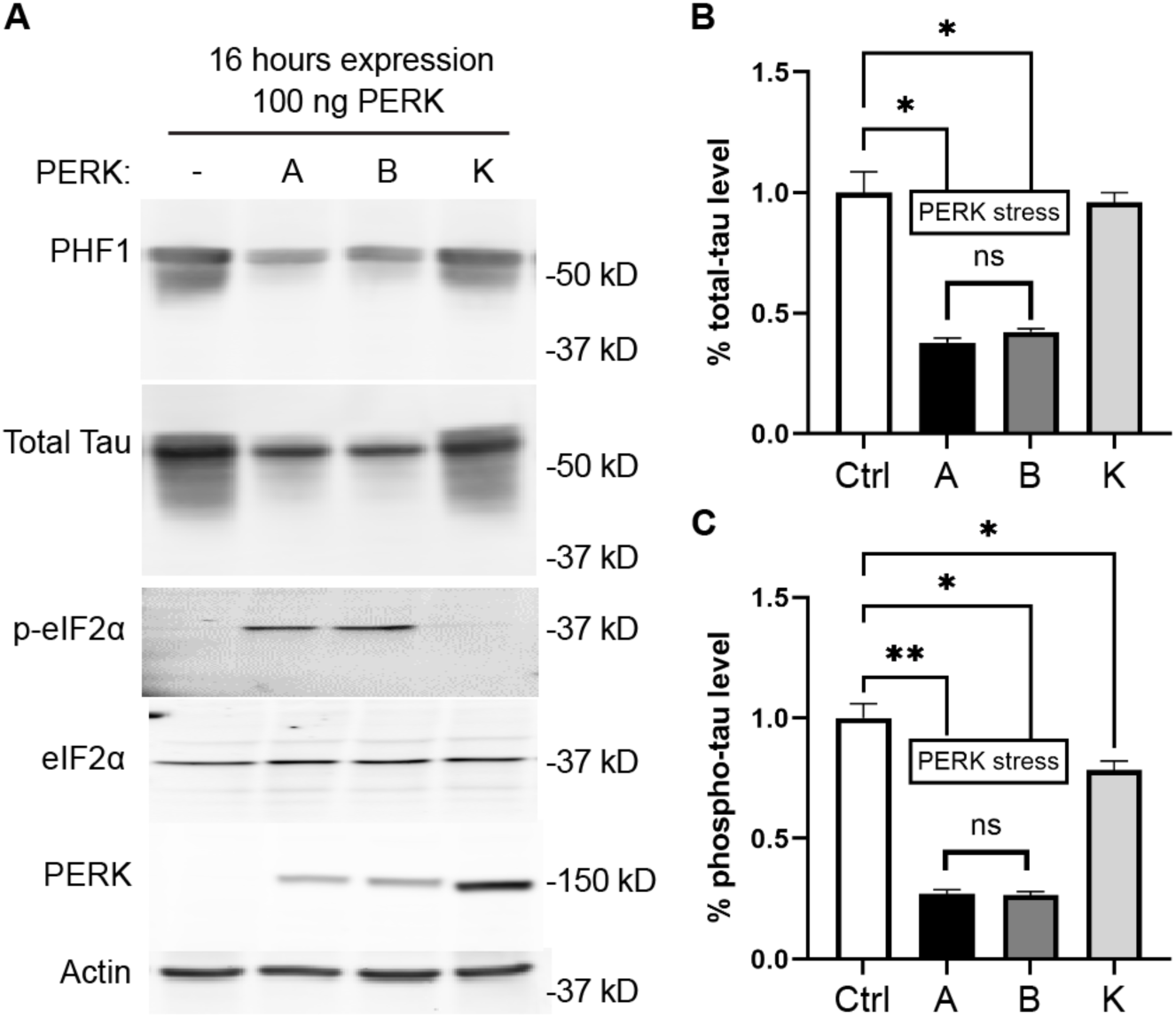
Short-term expression of PERK induces eIF2α phosphorylation without affecting tau phosphorylation. PERK KO HEK293T cells were co-transfected with 100 ng of PERK constructs (PERK-A, PERK-B; referred as PERK-stress), PERK-K, or an empty plasmid, and 1 μg of a plasmid expressing human wild type 0N4R tau. Cells were lysed 16 hours post-transfection, and proteins were analyzed by Western blot (A) and quantified by optical density analysis (B). Expression of PERK-A or PERK-B induced the phosphorylation of eIF2α compared to non-stress control or PERK K, indicating activation of the UPR but did not affect tau phosphorylation. Optical density data are presented as percentage mean ± SEM (n = 3; ns: not significant, *p < 0.05, **p < 0.01, one-way ANOVA, Tukey’s multiple comparisons).

We then transfected 50ng or 100ng of PERK expression plasmids for 48hrs (Fig. 4). We found that total PERK-A and PERK-B levels were increased in every condition; consistent with previous studies concluding that it is a hypomorph, PERK-B failed to produce significantly increased pPERK between 50ng and 100ng of plasmid (Fig. 4A-B). Interestingly, after 48hrs, neither 50ng nor 100ng conditions showed detectable eIF2α phosphorylation beyond controls (Fig. 4C).

**Figure 4.**
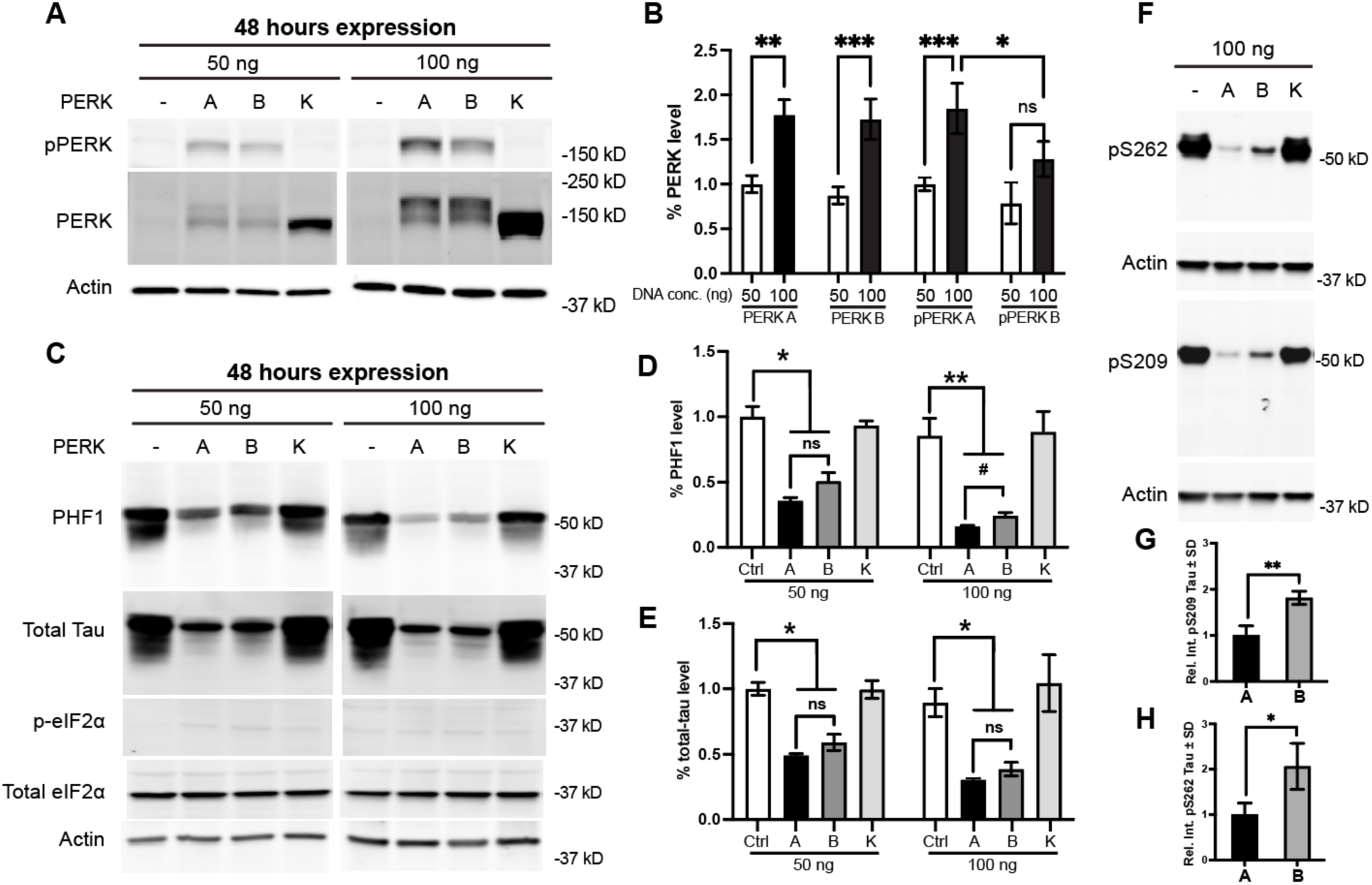
PERK-A expression reduces tauopathy-associated tau species more efficiently than PERK-B. PERK KO HEK293T cells were co-transfected with either 50 ng or 100 ng of PERK constructs (PERK-A, PERK-B; referred as PERK-stress), along with an empty plasmid and 1 μg of a plasmid expressing human 0N4R tau. Cells were lysed 48 hours post-transfection, and proteins were analyzed by Western blot and quantified by optical density. (A-B) Transfection with 100 ng of PERK-A or PERK-B resulted in higher PERK expression compared to 50 ng. PERK-A showed a significant increase in phosphorylation at Thr982 when comparing 100 ng vs. 50 ng, while PERK-B phosphorylation at Thr982 was not significantly higher at 100 ng and remained significantly lower than PERK-A at 100 ng. Optical density data are presented as percentage mean ± SEM (n = 3; ns: not significant, *p < 0.05, **p < 0.01, ***p < 0.001, two-way ANOVA, Tukey’s multiple comparisons). (C-E) Expression of PERK-A or PERK-B at both 50 ng and 100 ng reduced PHF1 and total tau levels compared to the non-stressed control. PERK-A showed lower PHF1 levels compared to PERK-B when transfected at 100 ng (*p < 0.05, ^#^p < 0.01, two-tailed paired t-test). Very low levels of phosphorylated eIF2α were detected under non-stress or PERK-stress conditions. (F-H) Western blot analysis of additional phospho-tau species (using the same samples from A) shows that PERK-A compared to PERK-B led to lower levels of tau pS262 and pS209. Optical density data are presented as percentage of the mean ± SEM (n = 3; *p < 0.05, **p < 0.01, two-tailed paired t-test).

Compared to controls, total tau levels were reduced following transfection with either PERK-A or PERK-B, with no significant difference in total tau levels between the two variants (Fig. 4C-E). Under conditions of higher PERK expression (100ng of cDNA for 48hrs), total tau levels remained unchanged; however, PERK-B was less effective than PERK-A at reducing PHF1 levels (Fig. 4C-D). Similar differences were observed in other disease-associated phospho-tau species, including pS262 and pS209 (Fig. 4F-H).

Another important feature of tauopathies is deposition of insoluble tau species. To test the impact of PERK haplotypes on tau aggregation, we co-transfected PERK-A, -B, or -K with a tau variant that contains two disease-associated mutations (P301L and S320F) and is highly prone to aggregation (termed 2x Tau) ^26^. We found that, compared to PERK-A, PERK-B less effectively reduced insoluble tau (Fig. 5A-B). Therefore, while expression of both PERK haplotypes associated with reduced pathological tau species, PERK-B less competently decreased pathological tau, suggesting that longer overexpression of PERK-B leads to accumulation of pathological tau species compared to PERK-A.

**Figure 5.**
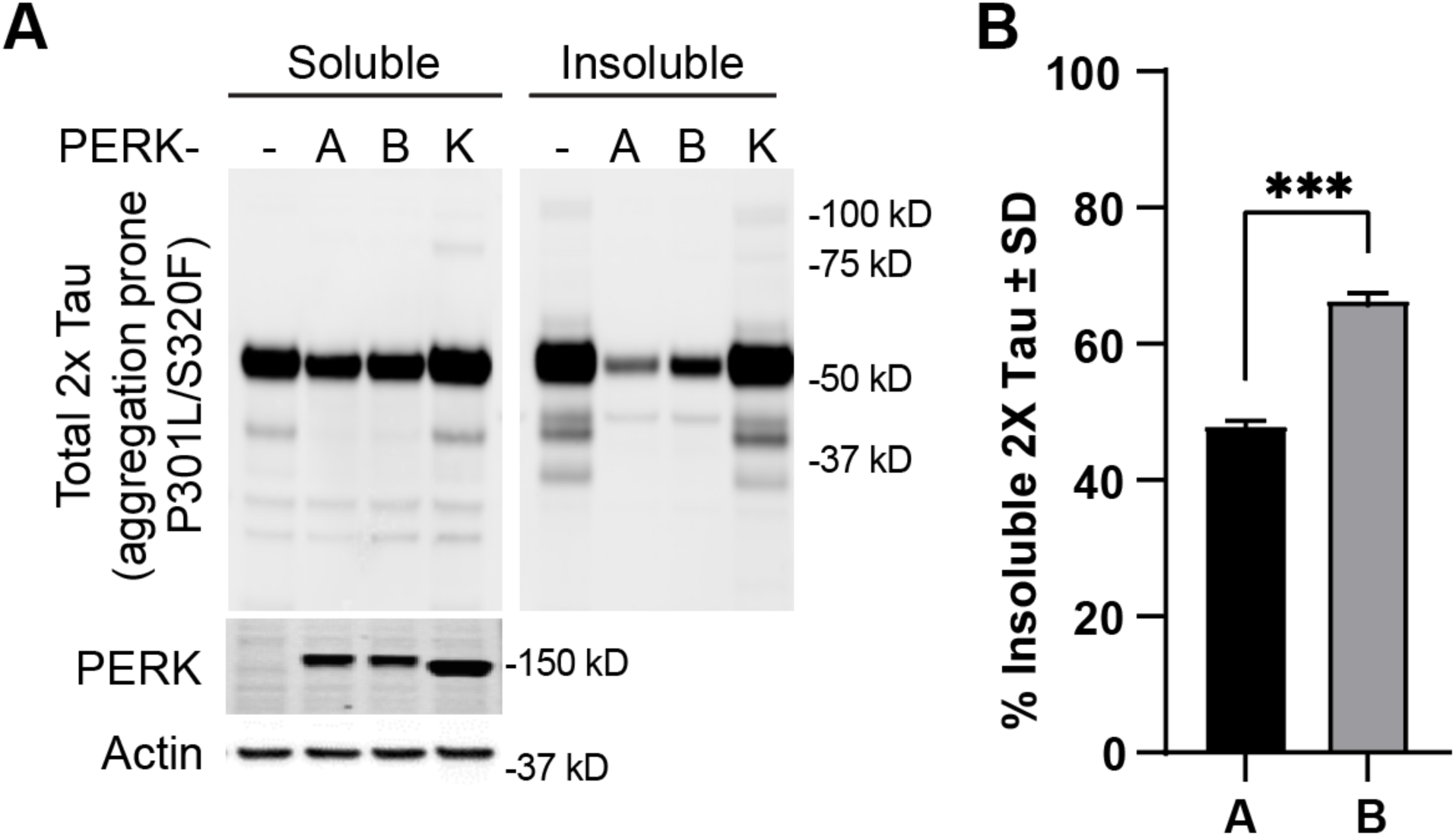
PERK-A expression reduces aggregation-prone 2x-tau (P301L/S320F) more efficiently than PERK-B. PERK KO HEK293T cells were co-transfected with 100 ng of PERK constructs (PERK-A, PERK-B, and PERK-K) or with an empty plasmid and 1 μg of 0N4R 2x-tau (P301L/S320F). Cells were lysed 48 hours post-transfection and subjected to ultracentrifugation to separate detergent-soluble and detergent-insoluble fractions. (A) Both fractions were analyzed by Western blot and quantified by optical density. (B) The PSP variant PERK-B resulted in a significantly higher percentage of aggregated (insoluble) 2x-tau compared to PERK-A. Optical density data are presented as the percentage of insoluble tau ± SEM (n = 3; *p < 0.05, **p < 0.01, two-tailed paired t-test).

### PERK-A and PERK-B transcriptomes are similar, but their translatomes are unique

To establish differences in PERK-A and -B that could explain how they differentially affect tau, we investigated changes in the transcriptome and the translatome. Our RNAseq experiments revealed stark differences resulting from PERK activation compared to control (Suppl. Fig. 1); however, there were no significant differences in the transcriptome between the haplotypes.

PERK’s canonical substrate, eIF2α, reduces overall translation, while promoting the synthesis of unique RNA sequences with upstream open reading frames. To identify whether the differences between PERK-A and PERK-B resided in their corresponding translatomes, we coupled SUnSET with puromycin immunoprecipitation and liquid chromatography/tandem mass spectrometry (LC-MS/MS).

We identified distinct translatomes for each sample: control, PERK-A, and PERK-B. We found that 46% of the PERK-A translatome was unique compared to control and PERK-B (Fig. 6). Meanwhile, only 2% of the PERK-B translatome was distinct from the other groups, suggesting that the risk imparted by PERK-B to develop PSP and AD is accounted within the 2% of nascently translated proteins. The PERK-A translatome exhibited an enrichment of proteins that suppress translation, as anticipated (Suppl. Fig. 2). While PERK-B-expressing cells shared some overall features of this translatome, the effect was significantly less pronounced compared to the profile observed in PERK-A cells. We identified and compared differentially expressed proteins (Suppl. Fig. 3) and found minor changes; PERK-B expression only reduced translation of the described transcripts. When listing the genes that were exclusively identified in each group, we found that only four proteins were uniquely identified in the PERK-B translatome: proteasome subunit alpha type-3 (PSMA3), alcohol dehydrogenase 5 (ADH5), distal-less homeobox 1 (DLX1), and neuroepithelial cell transforming gene 1 (NET1). Of these, only DLX1 was previously associated with PSP ^22^ (Table 1 and Fig. 6).

**Figure 6.**
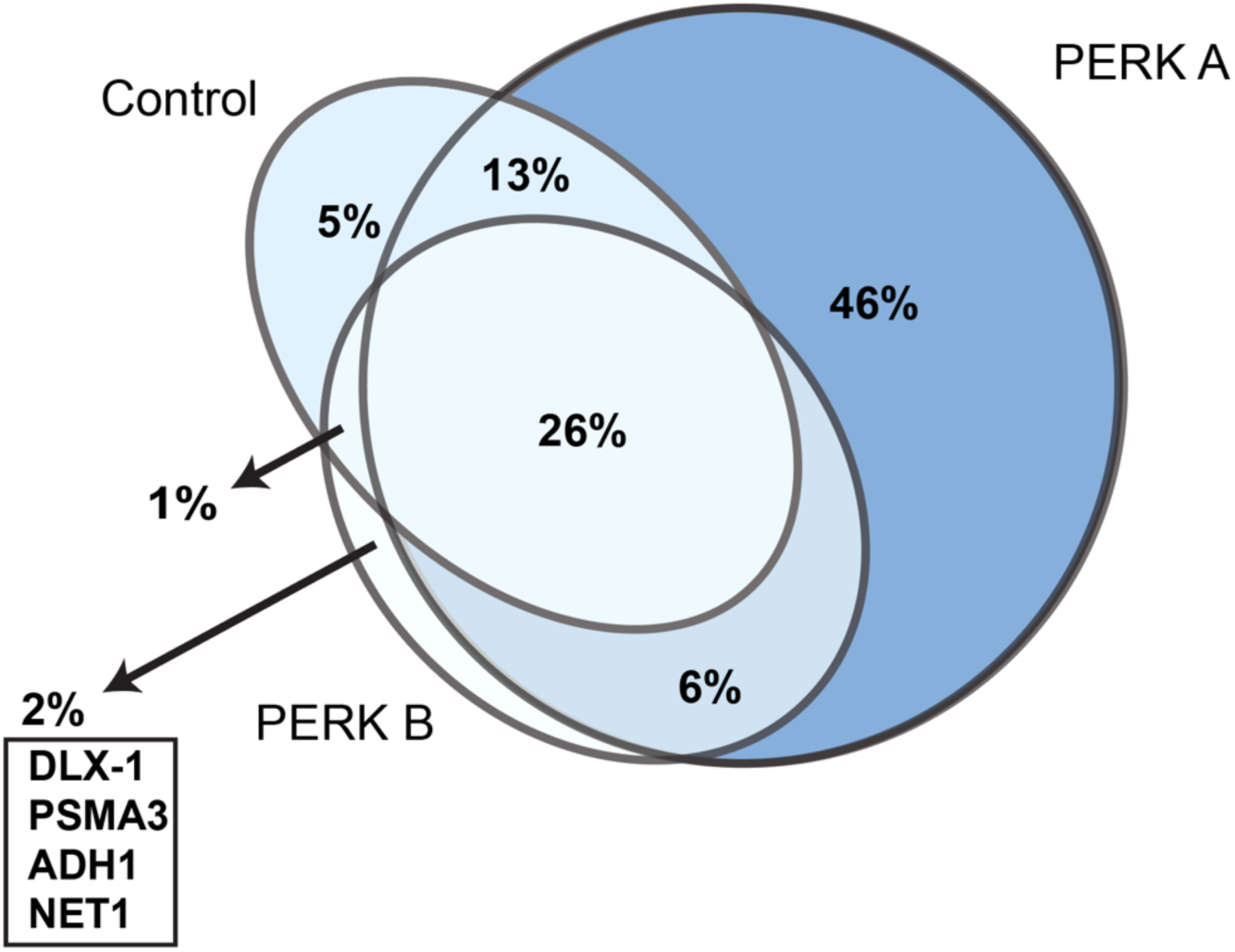
Expression of PERK-A or PERK-B alters global protein translation. PERK KO HEK293T cells were transfected with 400 ng of PERK constructs (PERK-A or PERK-B) along with an empty plasmid (control) in 10 cm dishes. At 48 hours post-transfection, cells were incubated with puromycin (10 µg/mL) for 1 hour to label nascent peptides. Protein lysates were collected and subjected to immunoprecipitation of puromycinylated peptides, followed by mass spectrometry analysis. Venn diagram shows the percentage of overlapping and unique peptides detected in each condition.

**Table 1:** Proteins identified in the PERK-B translatome but not in the PERK-A translatome.

| *Protein* | *-log10 (p-value)* | *Associated with PSP or AD* |
| --- | --- | --- |
| *ADH5* | 1.98 | No |
| *NET1* | 1.66 | No |
| *DLX1* | 1.54 | Weber et al. <sup>22</sup> |
| *PSMA3* | 1.51 | No |

To corroborate that DLX1 levels are altered in PSP, we measured changes in the accumulation of DLX1 in the soluble and insoluble fractions of human PSP striatum (Fig. 7). As expected, PHF1 tau levels were significantly increased in PSP brains (Fig. 7A). DLX1 levels were increased only in the detergent-insoluble fraction; the soluble fraction had decreased DLX1 (Fig. 7B-E), suggesting a role for DLX1 redistribution in pathological processes in tauopathies.

**Figure 7.**
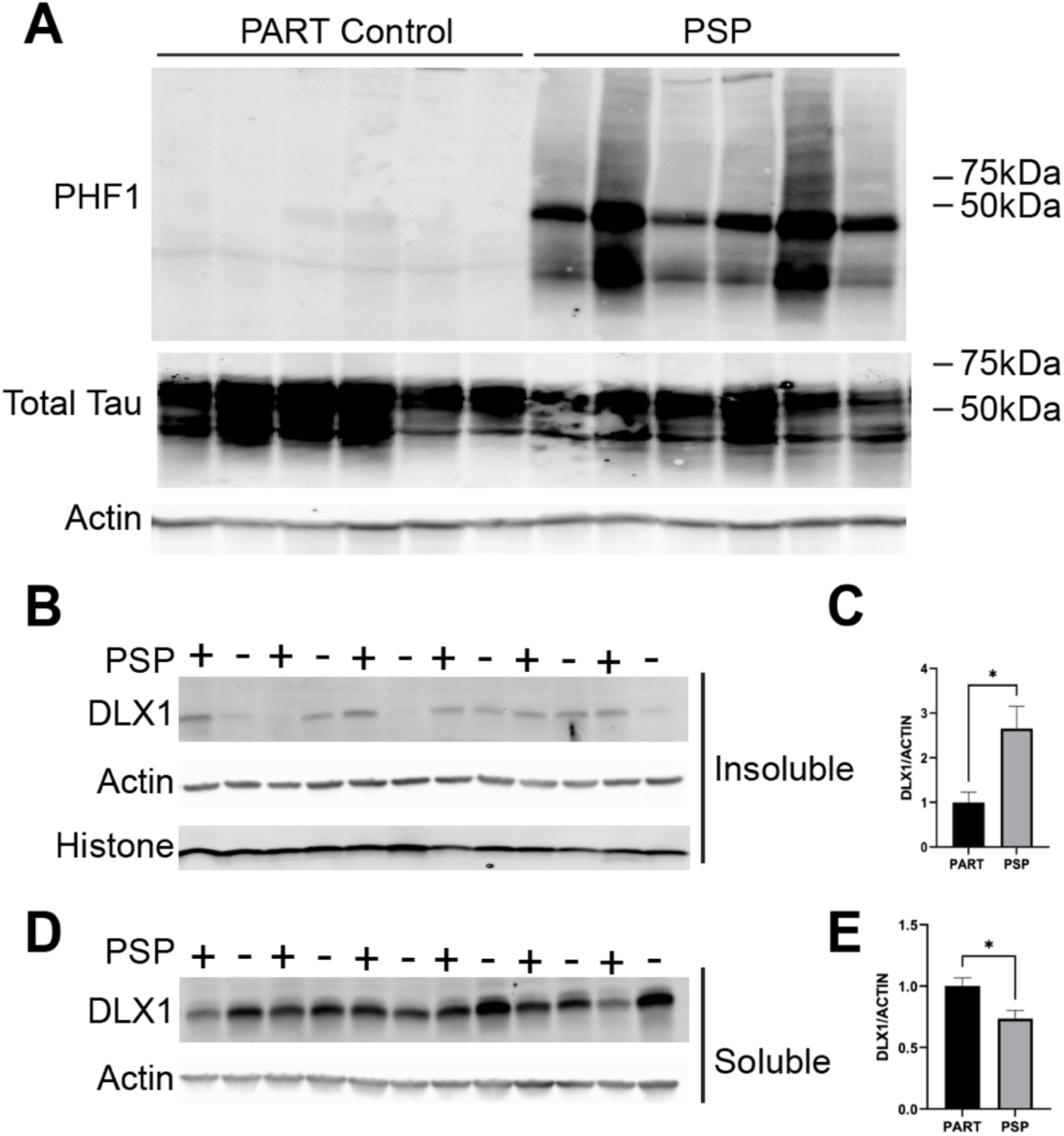
Human PSP brains present more insoluble DLX1. Striatum from PSP and early-stage Primary Age-Related Tauopathy (PART) human brains were processed to yield detergent-insoluble and detergent soluble fractions. (A) PSP striatum samples contain significantly more PHF1 tau compared to early-stage PART controls. DLX1 is significantly increased in the (B-C) detergent-insoluble fractions and significantly decreased in the (E-F) soluble fractions. Actin and histone protein was used as control. Optical density data are presented as the percentage of DLX1/actin ± SEM (n = 7-8; *p < 0.05, two-tailed paired t-test).

To investigate the effects of DLX1 *in vivo*, we used a Drosophila model of tauopathy in which tau overexpression in the eye leads to severe retinal disorganization and reduced eye size (Fig. 8). When the fly DLX1 homolog, distal-less (*Dll)*, was silenced in control flies, there was no change in eye morphology (Fig. 8A). Tau overexpressing flies showed significant alterations to eye morphology. However, *Dll* silencing in tau-overexpressing flies significantly rescued the eye defects (Fig. 8C-E), suggesting that DLX1 contributes to tau-mediated toxicity. Notably, *Dll* silencing had no effect on eye phenotypes induced by Aβ42 (Suppl. Fig 4), highlighting the specificity of the DLX1-tau effects over Aβ.

**Figure 8.**
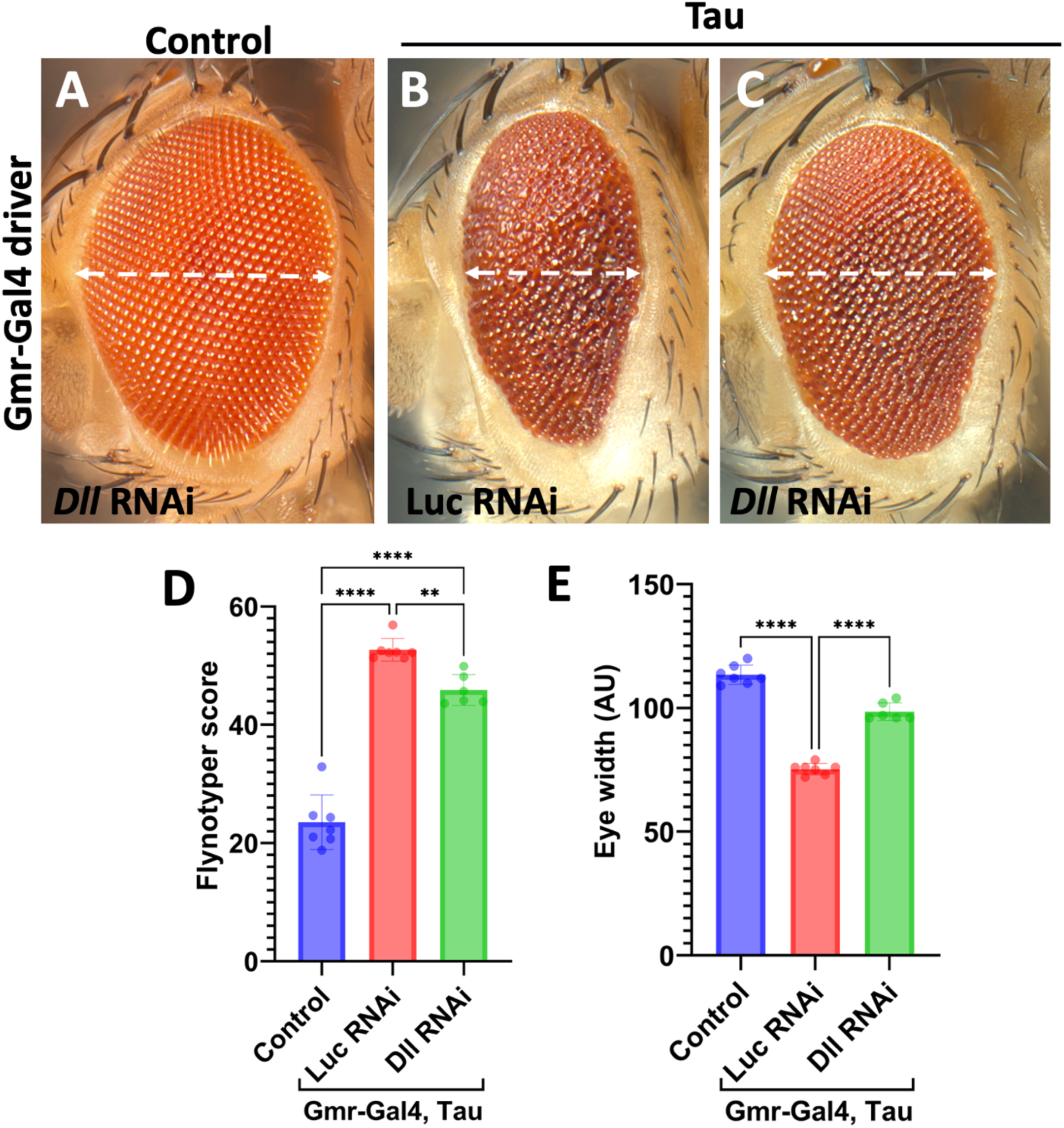
Distal-less/DLX1 silencing reduces tau toxicity *in vivo*. Representative eye images from Drosophila expressing either *Dll* RNAi alone (A), UAS-Tau with the innocuous Luciferase (Luc) RNAi control line (B), or UAS-Tau with *Dll* RNAi (C), under control of the eye-specific Gmr-Gal4 driver. Quantifications of the Flynotyper score (D) and eye width (E) are shown in arbitrary units (AU). The white dashed lines indicate maximum horizontal eye width measured with ImageJ (n≥7). Error bars represent SEM. Statistical significance was determined by one-way ANOVA with Tukey’s post-hoc test. **p < 0.01, ****p < 0.0001.”

## DISCUSSION

We describe a novel mammalian cell culture system that enables PERK expression at levels that induce the UPR without the addition of toxic chemical compounds such as thapsigargin or tunicamycin, and without inducing cell death. Using this system, we analyzed the impact of rare polymorphisms in the PERK gene that distinguish the A and B alleles on known PERK functions, including its broad suppression of translation. We established that canonical PERK functions were indistinguishable between the two variants. We observed no significant difference between PERK-A and -B in the activation of ATF4 and sXBbp1 transcription factors. PERK-A and -B showed similar capacities to inhibit cap-dependent translation. As previously described, PERK activation diminished phospho-tau levels, but PERK-B was slightly less effective than PERK A at reducing pTau epitopes we evaluated, confirming that this new model was consistent with previous systems ^12,27^. PERK B was also less effective in preventing the aggregation of mutant tau P301L/S320F. Utilizing this unique model in conjunction with puromycin-based proteomics, we uncovered a highly specific subset of proteins (comprising only 2% of the total PSP-associated proteome compared to controls) that are differentially regulated by PERK-A versus PERK-B. Among the identified genes, DLX1 stood out as a driver of toxicity in mammalian cell culture. We also show that DLX1 is increased in insoluble fractions from human PSP brains and that reducing the Drosophila homologue of DLX1 in fly eye mitigates degeneration caused by the over-expression of WT human tau. Thus, our data link, for the first time, a toxic PERK-DLX1 pathway with tauopathy in human disease.

Increased activation of PERK correlates with tau pathology in PSP and AD brains^15^. We confirmed that beyond this positive correlation, PERK haplotypes differentially regulate tau and phospho-tau protein levels. However, Stutzbach and colleagues showed that immunoreactivity for active PERK and tau do not overlap in all cells. Additionally, increased pPERK has been detected in neurons absent of neurofibrillary tangles in brains from AD cases^17^. Therefore, it is important to determine whether expression of PERK haplotypes is cell specific, which is likely given results from previous work showing that astrocytic and neuronal PERK promote different UPR outcomes ^28,29^. Nonetheless, our data suggest that when tau is present in cells with PERK-B, the levels of insoluble tau will be elevated.

The PERK-B haplotype confers higher sensitivity to ER stress, inefficiently regulates eIF2α, has limited self-activation, and promotes accumulation of pathological tau ^9,15,30^. All these are mechanisms by which PERK-B carriers may have a higher risk for neurodegenerative diseases^15^. However, until now, these PERK mechanisms did not link the transcription factor DLX1 in a proteotoxic pathway with tau. We identified a highly specific PERK-B translatome (Table 1) that corresponds to only 2% of all proteins unique to PERK-B compared to control (non-stressed) and PERK-A. DLX1 was the only protein identified that had been previously associated with PSP. Additional work is needed to evaluate whether the other PERK-B-associated proteins may impact tau directly and/or play a role in PSP. Specific efforts could focus on whether PSMA3, ADH5, and NET1 also modulate pathological tau accumulation, contribute to indirect pathways (like PKR protein), are distributed in main brain regions affected in tauopathies, or exert control over protein regulation (via the proteasome, autophagy, and enzymatic activity). Our study identifies DLX1 as a key protein that influences tau toxicity in multiple systems, and it is strongly supported by previous work from Weber et al. genetically implicating DLX1 in PSP ^22^. Further research is required to address the precise mechanisms through which DLX1 modulates tau phosphorylation and aggregation.

The inefficiency of PERK-B in controlling tau levels can be explained by its limited self-phosphorylation and reduced kinase activity which can lead to weak eIF2α activation after stress induction ^9^. Because tau levels increase with age correlating with declined proteostasis^31^, this delay and partial PERK-B activation over time can be detrimental for cells to cope with age-related cellular stress. Certainly, unregulated proteins synthesis has the potential to create dyshomeostasis dyshomeostasis possibly contributing to cell death caused by misfolded tau.

Our findings highlight the significant role that PERK haplotype variations play in shaping the cellular proteome in the context of PSP and draw greater attention to the importance of translational regulation pathways in neurodegenerative diseases. This disease-associated translatome had not been previously associated with PSP, underscoring the novelty of these results and the opportunity it opens to unravel disease mechanisms.

## Material and methods

### Antibodies

PERK was detected with D11A8 Rabbit (Cell Signaling #5683) or Anti-FLAG M2 (Sigma, F1804) antibodies. Tau was detected with rabbit antibody 3026 that was raised against recombinant 0N3R human^32^. β-Actin was detected with AC-15 (Santa Cruz, sc-69879). Anti-Puromycin antibody (clone 12D10) was purchased from EMD Millipore (#MABE343) Secondary antibodies detection for Western Blot IRDye 800CW (#926-32212) was purchased from LI-COR Biosciences and Alexa Fluor 680 rabbit(#A-21109) was purchased from Thermo Scientific.

### Plasmids

All plasmids used in this study (listed in Suppl. Table 1) were approved by the Environmental Health and Safety Office at the University of Florida. The human PERK haplotype A plasmid was obtained from Origene (#RC214993). Mutations corresponding to PERK haplotype B (S136C, R166G, and S704A), as previously described ^9^, as well as the K622R mutation, were introduced into PERK-A by PCR-based mutagenesis and confirmed by DNA sequencing. The plasmids encoding ATF4-mScarlet-NLS (Addgene #115969) and pLVX-XBP1-mNeonGreen-NLS (Addgene #115968) were gifts from Dr. David Andrews ^33^. Human 0N4R tau was cloned into pcDNA3.1. The double mutant tau construct (“2X-TAU”, 0N4R P301L/S320F) was digested with *BlpI* and *NheI*, and the tau fragment was subcloned into the 0N4R tau backbone. An alternative open reading frame (AltORF) strategy previously described^34^ was used to enable expression of a second protein in the +3 reading frame of EGFP. To generate this construct, mApple fluorescent protein was subcloned into pENTRY-EGFP using the *XhoI* restriction site, positioned C-terminally after the EGFP stop codon in the +3 reading frame. The correct insertion and reading frame of mApple were verified by DNA sequencing.

### Cell lines and generation of HEK293T PERK knock-out cells

HEK293T cells were grown in DMEM supplemented with Corning 10% fetal bovine serum (GE) and 100 units/mL penicillin, 100 µg/mL streptomycin (Life Technologies). PERK knockout HEK293T cell line was generated with CRISPR/Cas9 genome-editing^35^. A RNA guide aiming exon 3 of PERK (sense CACCGTTTCCATGCTTTCACGGTCT and antisense AAACAGACCGTGAAAGCATGGAAAC) was subcloned in pSpCas9(BB)-2A-GFP (gift from Feng Zhang, Addgene #48138). Plasmid expressing the cloned gRNA was confirmed by sequencing and transfecting in HEK293T. Transfected cells were plated on 96-well plates and single clones were analyzed for endogenous PERK expression level by Western blotting. Genomic DNA from colonies of interest was extracted for PCR analysis with the following primer (forward primer GCTCTTAATTACTATTATTC, reverse primer GATTGCAAAATAATGTATAA). PCR products were cloned in TOPO-TA (Thermo Fisher, #4500781). Cells were transiently transfected using calcium phosphate DNA-precipitation method^36^. We used pcDNA3.1 as DNA carrier for small amount of transfected PERK.

### Western Blotting

Protein extracts were mixed 1:1 with 4X protein loading buffer (31.25 mM Tris-HCl pH 7.5, 2% LDS, 10% glycerol, 1.5% β-mercaptoethanol, and Orange G), then electrophoretically separated on 12–4% Bis–Tris precast gels or 12–4% Tris–glycine gels (Bio-Rad) as previously described ^37,38^. Proteins were electrophoretically transferred onto Immobilon-PSQ PVDF membranes (Millipore Sigma, #ISEQ00010) and subsequently blocked in Tris-buffered saline (TBS) containing 0.5% casein for 3 hours at room temperature. Primary antibodies were diluted in TBS with 0.2% Tween-20 (TBS-T) and incubated for 1 hour at room temperature or overnight at 4°C. Membranes were washed three times (5 minutes each) in TBS-T, followed by a 1-hour incubation with secondary antibodies diluted in TBS-T. After an additional three, 5-minute washes in TBS-T, membranes were imaged using the Odyssey Infrared Imaging System (LI-COR Inc.). Biochemical cellular fractionation of 2X-TAU was performed as previously described ^39^.

### Puromycin assay

Transfected cells were incubated with puromycin (10 µg/mL; 18.36 µM) for 1 hour to label nascent peptides as previously described ^37,40^. Following incubation, cells were lysed in RIPA buffer supplemented with EDTA-free protease inhibitor cocktail (Sigma) and 1 mM PMSF (Sigma). Lysates were sonicated and then centrifuged at 13,000 rpm for 15 minutes at 4°C. The resulting supernatants were collected, and protein concentrations were determined before analysis by Western blotting.

### RNA extraction and RT-PCR

Total RNA was extracted from transfected cells using TRIzol reagent (Invitrogen), following the manufacturer’s instructions. Reverse transcription (RT) was performed using the High-Capacity cDNA Reverse Transcription Kit (ThermoFisher Scientific), with random primers for general transcript analysis and oligo(dT) primers for specific detection of XBP1 mRNA splicing. Subsequent PCR amplification was carried out using Taq 2X Master Mix (New England Biolabs, #M0270) and gene-specific primers. Tau mRNA was amplified using a forward primer targeting an internal tau sequence (5’-GCAAATAGTCTACAAACCAGTTG-3’) and a reverse primer (5’-TAGAAGGCACAGTCGAGG-3’) complementary to the BGH polyadenylation sequence in the plasmid, ensuring detection of only the exogenous tau transcript. XBP1 was amplified with primers: forward (5’-AGAACCAGGAGTTAAGACAGC-3’) and reverse (5’-AGTCAATACCGCCAGAATCC-3’). GAPDH was amplified as a control using validated primers from Origene (Catalog #HP205798).

### RNA Sequencing and Data Analysis

PERK constructs were transfected into PERK knockout (KO) HEK293T cells, and total RNA was extracted 48 hours post-transfection using TRIzol reagent (Invitrogen), following the manufacturer’s protocol. Total RNA sequencing and bioinformatic analysis were performed by LC Sciences (Houston, TX, USA). Differential gene expression analysis was conducted, and significantly regulated genes were identified using a false discovery rate (FDR) threshold of q < 0.05. The resulting dataset was analyzed in R, and heatmaps were generated using the pheatmap package (Kolde R, 2018. pheatmap: Pretty Heatmaps. R package version 1.0.12, https://github.com/raivokolde/pheatmap) to visualize expression patterns of differentially expressed genes.

### Puromycin Proteomics

As previously performed ^24,38,40^, transfected cells expressing PERK constructs were incubated with puromycin (10 µg/mL; 18.36 µM) for 1 hour, except for the negative control group, which was used to distinguish specific puromycinylated peptides from background or non-specific proteins bound to the antibody–Dynabead complex.

Following treatment, cells were washed with PBS and lysed in a buffer containing 150 mM KCl, 10 mM MgCl₂, 5 mM HEPES (pH 7.4), and 1% Igepal CA-630, supplemented with EDTA-free protease inhibitor cocktail (Sigma) and 1 mM PMSF (Sigma). In parallel, 5 µg of anti-puromycin antibody was crosslinked to Dynabeads Protein G (NEB, #S1430S) following the manufacturer’s protocol and as previously described ^40^. Cell lysates were clarified by centrifugation at 16,000 × g for 15 minutes at 4°C, then incubated with the antibody-crosslinked Dynabeads for 16 hours at 4°C on a rotator. After incubation, the immunoprecipitated complexes were washed three times with PBS containing 0.2% Tween-20, followed by additional washes with a larger volume of PBS to remove residual detergent. The bound puromycinylated nascent peptides were subsequently analyzed by mass spectrometry at Emory University.

### Mass Spectrometry Data Analysis

Raw mass spectrometry data were processed as previously described with minor adjustments ^37,41^. The resulting MaxQuant output files were imported into Perseus for initial filtering. Data were then exported to Excel for further statistical analysis in R, due to the large dataset size. Peptides were initially filtered to retain only those with at least three positive intensity values in one experimental group. For entries with two positive and one missing or null value, a permutation step was applied by replacing the missing value with the average of the two available values. Peptide detection was assessed relative to the negative control group. A peptide was considered positively detected if it showed a measurable intensity signal while the corresponding control value was zero. Conversely, peptides were classified as not detected if their average intensity in the experimental group was equal to or lower than the average intensity in the negative control. Statistical comparisons between each experimental group and the negative control were performed using one-tailed t-tests, with test selection based on variance equality (Student’s t-test for equal variance, Welch’s t-test for unequal variance).

Peptides with a significantly higher mean intensity (p < 0.05) compared to the negative control were classified as positively detected. Subsequently, two-tailed t-tests were conducted to assess differences between experimental groups.

### Gene Ontology and Data Visualization

Gene Ontology (GO) enrichment analysis was performed using g:Profiler ^42^ to identify biological processes associated with peptides uniquely detected in each experimental group. Venn diagrams were generated in R using the ggVennDiagram package (Gao C, Dusa A, 2024. ggVennDiagram: A ’ggplot2’ Implement of Venn Diagram. R package version 1.5.3. Available at: https://gaospecial.github.io/ggVennDiagram/, https://github.com/gaospecial/ggVennDiagram). Heatmaps were created using the pheatmap package (Kolde R, 2018. pheatmap: Pretty Heatmaps. R package version 1.0.12. Available at: https://github.com/raivokolde/pheatmap) to visualize Z-scores of differentially detected peptides.

### Drosophila Genetics and Eye Analysis

Drosophila stocks for the eye-specific driver Gmr-Gal4 (B-1104) and the UAS-Luciferase RNAi control (B-31603) were obtained from the Bloomington Drosophila Stock Center in Indiana, whereas the *Dll* RNAi line (v101750) was purchased from the Vienna Drosophila Resource Center. UAS transgenic flies expressing the longest human wild-type tau isoform (2N4R) were previously described as well as flies expressing human Aβ42 fused to a secretory signal peptide ^43,44^ . For the experiments, we collected virgins from recombinant stocks carrying Gmr-Gal4, UAS-Aβ42/Cyo and Gmr-Gal4, UAS-Tau/Cyo, and crossed them with males from UAS lines expressing Luciferase RNAi or *Dll* RNAi transgenes. All the crosses were cultured on pre-made molasses-based food from Archon Scientific Inc. at 26 °C. Eye phenotypes from progenies were documented using a Leica Z16 APO macroscope as described ^43^. Quantifications were performed using NIH ImageJ’s measure tool and Flynotyper plugin to calculate differences in size and organization of the fly eyes, respectively.

### Statistical analysis

Statistics were performed with GraphPad Prism (USA) version 9.4. All data are expressed as the mean ± SEM or ± SD as described.

## Acknowledgments

S.P. is the Charlotte and Howard Zimmerman Rising Star Professor with the Norman Fixel Institute for Neurological Diseases.

## Author contributions

Following the ICJME COPE guidelines, CL, JFA, TΝG, DB, DC, MJL, and DRL contributed to the conception and design of all or parts of the study. Data were acquired and analyzed by CL, JFA, ST, KM, PB, NS, DRR, PB, DC, JP, and SP. CL, JFA, JK3, MCM, SR, DB, TEG, TΝG, and DRL performed or contributed to data interpretation. All authors contributed to preparation of the manuscript.

## Data Availability Section

The datasets and computer code produced in this study are available upon request.

## Funding and additional information

This work was in part supported by the National Institutes of Health/National Institute on Aging (JFA R01AG075900-01, JFA R01AG074584-02, JFA R21AG093972-01, DRL R01AG077534) National Institute for Neurological Disorders and Stroke (JFA R01NS091329), the Alzheimer’s Association (JFA AARG-D-21-847204), and the Rotary Club (JFA Coins for Alzheimer’s Research Trust). The UF Neuromedicine Human Brain and Tissue Bank is supported by the Center for Translational Research in Neurodegenerative Diseases, the Evelyn F. and William L. McKnight Brain Institute and the 1Florida ADRC (NIH/NIA P30AG066506).

---

**Supplementary Figure 1.**
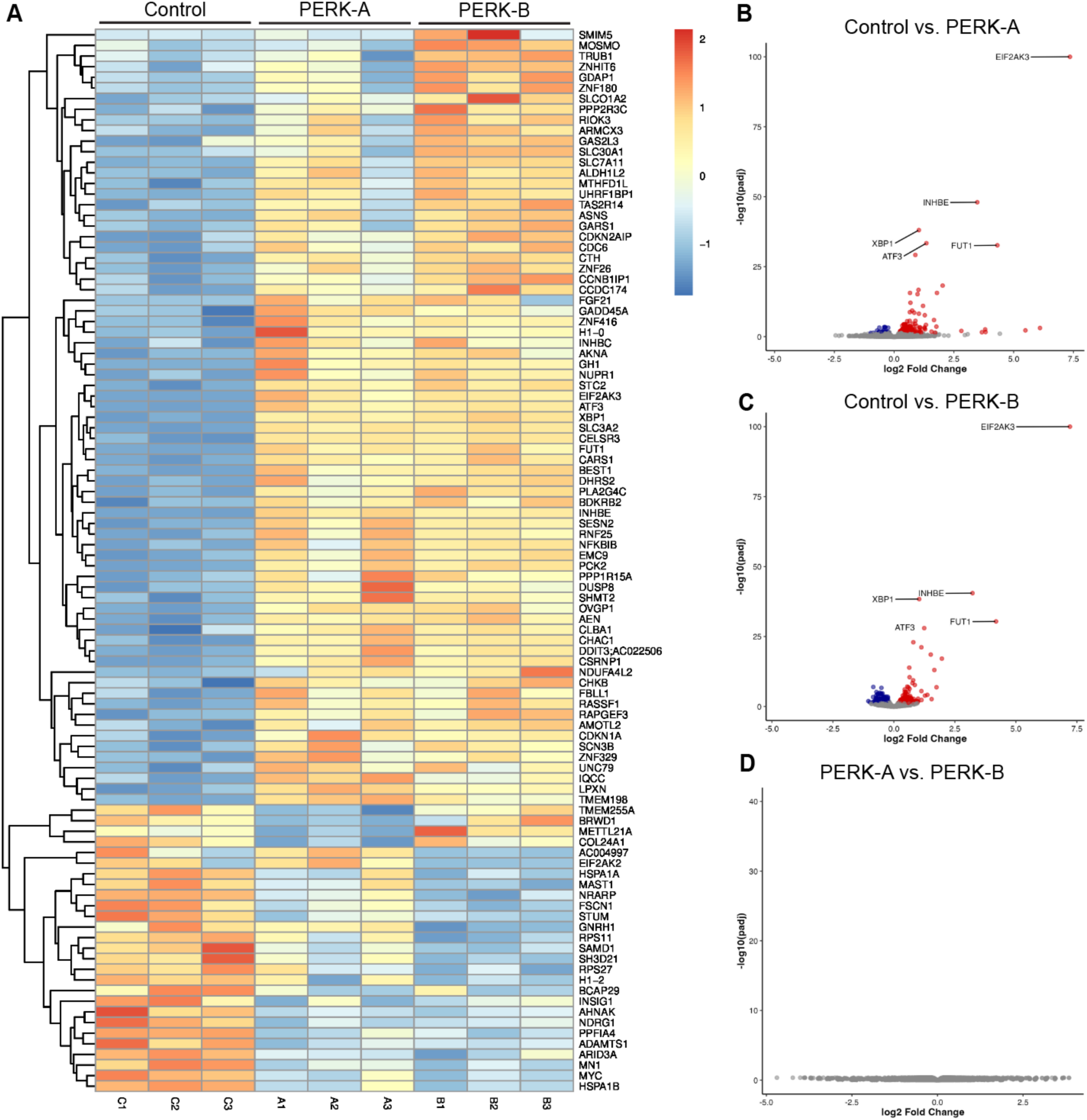
PERK-A and PERK-B expression similarly affect gene transcription profiles. PERK KO HEK293T cells were transfected with 100 ng of PERK constructs (PERK-A or PERK-B) along with an empty plasmid as a carrier. Cells were harvested 48 hours post-transfection for RNA isolation and transcriptomic analysis. (A) Clustered heatmap of differentially expressed genes (rows) across experimental conditions (columns: C1-C3 = Control replicates, A1-A3 = PERK-A replicates, B1-B3 = PERK-B replicates). The color scale represents the z-score normalized expression values. (B) Volcano plot showing differentially expressed genes in Control vs. PERK-A. The x-axis represents log2 fold change, and the y-axis represents -log10(adjusted p-value). Select genes of interest are labeled. (C) Volcano plot showing differentially expressed genes in Control vs. PERK-B. (D) Volcano plot showing that PERK-A vs. PERK-B genes are not differentially expressed.

**Supplementary Figure 2.**
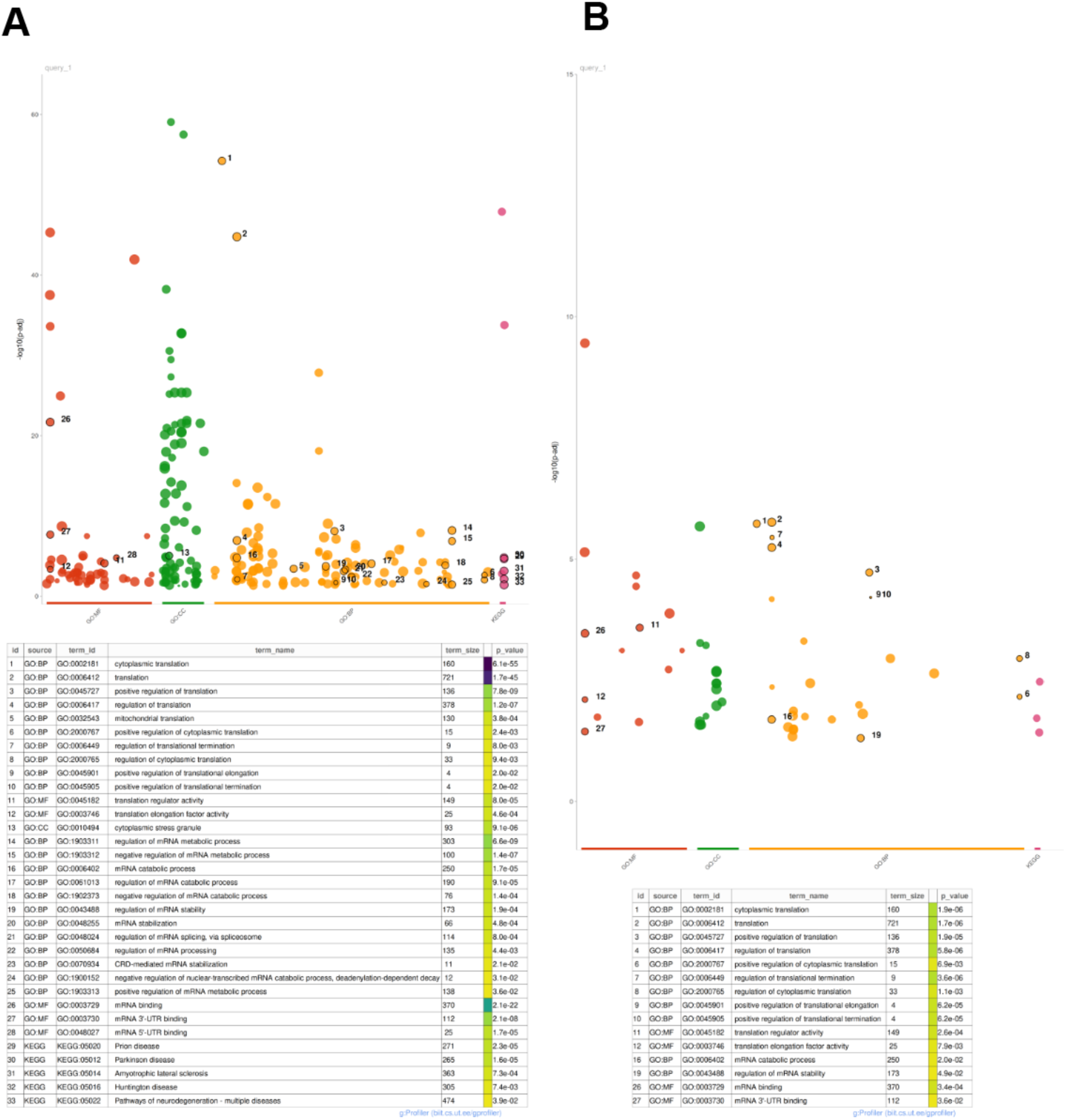
Gene Ontology (GO) analysis of peptides uniquely detected in PERK-A and PERK-B conditions. Peptides uniquely identified in PERK-A and PERK-B conditions from the mass spectrometry dataset were analyzed using the g:Profiler tool for Gene Ontology (GO) enrichment. Selected GO terms are visualized using Ghost plots, presented in (A) for PERK-A and (B) for PERK-B, highlighting functional categories enriched under each condition.

**Supplementary Figure 3.**
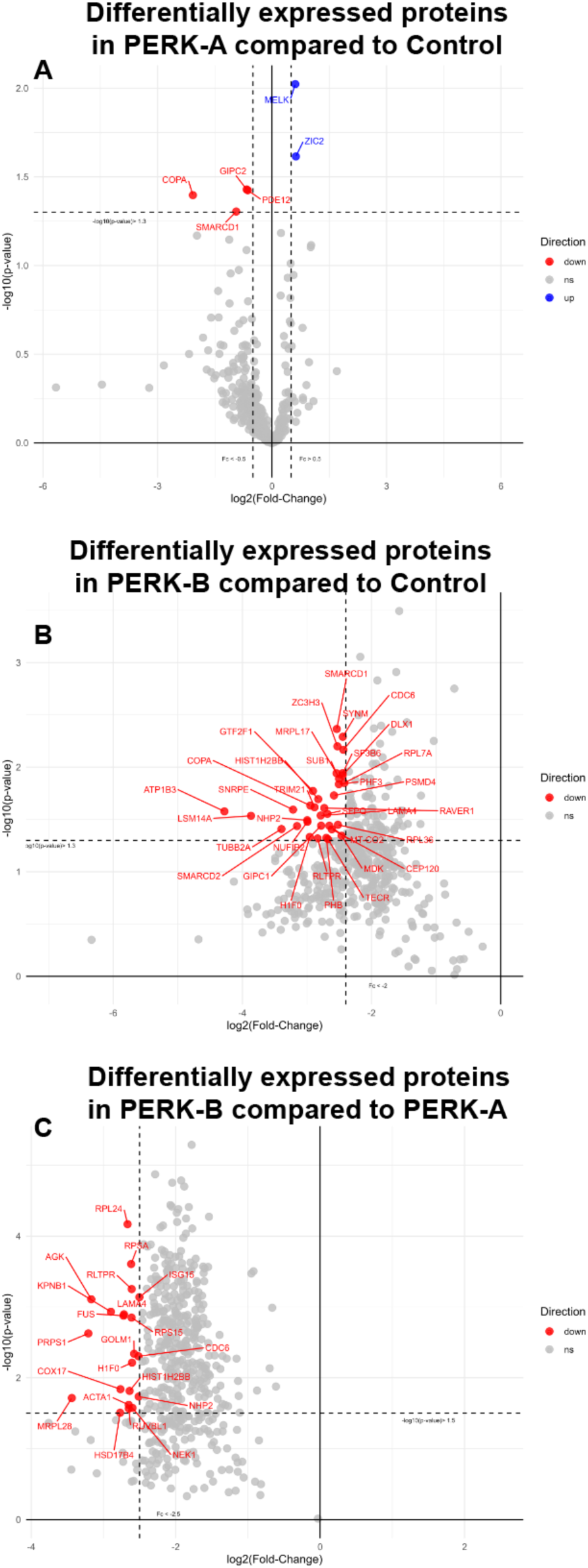
Volcano plots of peptides uniquely detected in PERK-A, PERK-B, and control conditions. Peptides uniquely identified from the puromycin mass spectrometry datasets were compared between (A) PERK-A and control, (B) PERK-B and control, and (C) PERK-A and PERK-B. PERK-B expression primarily promoted reduced protein synthesis when compared to PERK-A or control.

**Supplementary Figure 4.**
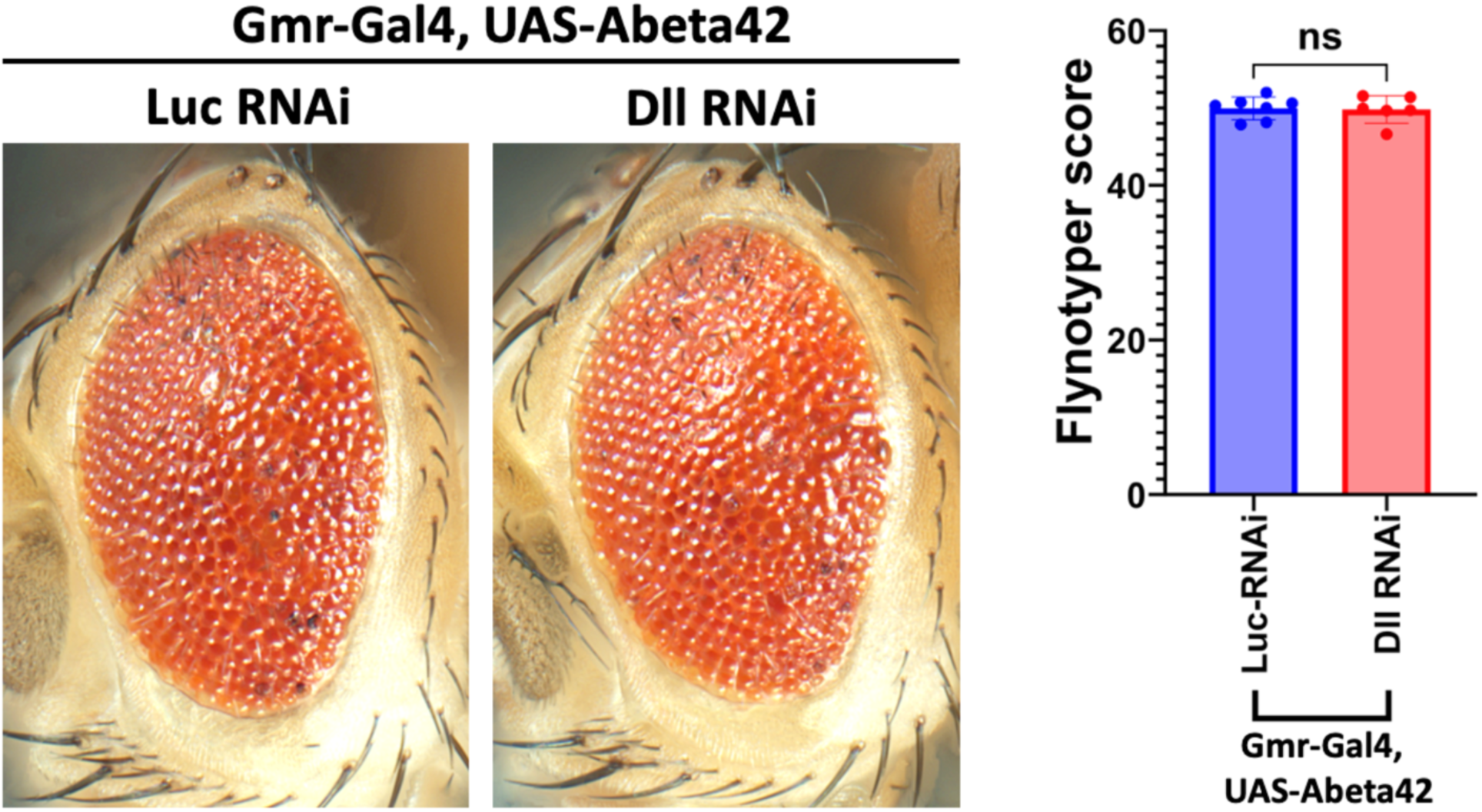
Distal-less/DLX1 silencing does not change eye morphology in a model of Aβ 42 overexpression. Representative images of adult Drosophila eyes co-expressing the indicated transgenes under control of the eye-specific Gmr-Gal4 driver. Flies co-expressing Aβ42 and the Luciferase (Luc) RNAi control show similar phenotypes to flies co-expressing Aβ42 and *Dll* RNAi, indicating no modification of Αβ42 toxicity by *Dll* silencing.

## Supporting information

This article contains supporting information.

**Supplemental Table 1:** Proteins identified in the PERK-A translatome but not in the PERK-B translatome.

| *Protein* | *-log10 (p-value)* |
| --- | --- |
| *CC2DB* | 2.98 |
| *GIPC2* | 1.98 |
| *SF3A2* | 1.87 |
| *IPO5* | 1.87 |
| *ATP2A2* | 1.83 |
| *PDE4DIP* | 1.81 |
| *RBM15B* | 1.70 |
| *SP110* | 1.61 |
| *DNAJC7* | 1.56 |
| *AKR1B1* | 1.44 |

**Supplemental Table 2:** Proteins identified in the PERK-A translatome but not in the control translatome.

| *Protein* | *-log10 (p-value)* |
| --- | --- |
| *MRPL10* | 5.72 |
| *MSX2* | 4.58 |
| *APOB* | 4.55 |
| *SNRNP70* | 4.39 |
| *THBS1* | 4.24 |
| *E2F7* | 3.23 |
| *NOP56* | 1.39 |
| *MRPL10* | 5.72 |
| *MSX2* | 4.58 |
| *APOB* | 4.55 |

